# The metatranscriptomic response of the wheat rhizosphere to drought varies with growth stages

**DOI:** 10.1101/2025.11.26.690733

**Authors:** Pranav Mukund Pande, Julien Tremblay, Marc St-Arnaud, Etienne Yergeau

## Abstract

Microbes can help plant sustain abiotic stresses, such as drought. Plant-microbe interactions are, however, dynamic and the timing of the stress will affect both partners, directly and indirectly. Here, we hypothesize that the effect of drought stress on the wheat rhizosphere microbiome would change between key growth stages. We grew wheat in pots and reduced soil water content for two weeks at stem elongation, booting, or heading. We then sampled the rhizosphere soil and sequenced its metatranscriptome. The timing of the drought strongly affected the transcriptional response of the microbes, but few differentially abundant transcripts were shared across all stages. Some common patterns were however observed at higher taxonomical or functional levels. Drought also affected the normal succession across wheat growth stages. Many of the differentially abundant transcripts, taxa and functions between growth stages of the control plants were not significant anymore for plants that experienced drought. Our results suggest that the timing of the drought event is paramount to the microbial rhizosphere communities, and that it could explain the heightened sensitivity of younger plants to stresses.

## Introduction

Drought can – depending on the growth stages – kill crops, severely reduce yields and grain quality [1], or have no effect. However, it is not clear if this is due to direct effects on the plant or indirectly through changes in beneficial plant-microbe interactions. A recent metatranscriptomic study from our group supports the indirect route. First, the microbiota – and not the plant – was the main responder to decreasing soil water content [2]. Second, this decrease in water content resulted in higher abundance of microbial transcripts that might be beneficial to the plant and microbes [2]. For instance, one microbial strategy to adapt to the decreased matric potential in dry soil is to accumulate osmolytes [3], and accordingly, we observed more microbial transcripts related to osmolytes in soils with low water content [2]. This could also be beneficial for the plant as microbes can exude osmolytes in the plant environment [4, 5]. Plant tolerance to drought- or salinity-related stresses can indeed be improved by different strains of *Actinobacteria* and *Proteobacteria* [6–9] and by fungal endophytes [10–12]. These beneficial microbial taxa and associated functions will, however, vary with plant growth stages, which could partly explain the varying sensitivity of crops to drought. This interaction between drought and growth stage is not fully understood, as most of the above-mentioned studies focused on a single sampling.

Time and water stress directly, indirectly, and interactively affects microbial communities. First, microorganisms respond directly to soil water availability [13–15], which contributes to seasonal changes in microbial communities [16, 17]. Second, water-stress induced shifts in root exudation [18, 19] can result in indirect effect of water stress on plant-associated microorganisms. Third, time can indirectly affect plant-associated microbial communities as root exudation change with plant growth stage [20, 21], shifting the microbial communities Finally, indirect and direct effects of time and water stress interact to influence the microbial communities. For instance, wheat associated microbial communities were altered by historical and current water stress in interaction with plant growth stage, plant genotype, and plant compartment [13–15]. As most studies look at single factors in isolation, it is difficult to predict the impact of the timing of drought on the microbial response and its feedback to crops, even though this knowledge would be crucial to devise microbial approaches to help crops adapt to water stress.

Global change will result in decreased soil moisture and increased drought events [22, 23]. Drought can reduce wheat yield by up to 50% [24, 25]. We think that if their response to drought was better understood, microorganisms could help abate these losses. As a first step, we need to understand how microbial communities will respond to water stress at different wheat growth stages. To answer this question, we grew wheat plants in pots without watering them for two weeks at three distinct growth stages (stem elongation, booting, and heading). At tillering, booting, stem elongation, heading and flowering, we uprooted plants, sampled their rhizosphere soil, extracted the RNA and sequenced it. We hypothesized that the effect of drought stress on the wheat rhizosphere microbiome would vary between key growth stages, with the microbiome of younger plants being more affected.

## Material and methods

### Experimental design and sampling

To test our hypothesis, we set-up a greenhouse pot experiment (Fig. 1). Each 1L pot was filled with 1.5 kg of dry soil. The soil was collected from our experimental field at the Institut national de la recherche scientifique (Laval, QC, Canada). This soil is naturally subjected to dry-rewet cycles during the summer [17]. All the soil was air dried, sieved with a 2 mm mesh sieve and homogenized. We then sowed in each pot 8 seeds of *Triticum aestivum* cv. AC Nass (Semences Nicolet, Nicolet, QC, Canada), a drought-sensitive cultivar. After germination, the plants were thinned to 5 per pot. Until the stem elongation stage, all the pots were maintained at 50% SWHC (soil water holding capacity). Drought was imposed for two weeks by dropping to 8% SWHC at the beginning of stem elongation, booting and heading growth stages, resulting in 3 treatments plus a control that was kept at 50% SWHC throughout the experiment (Fig. 1). We wanted to impose drought during the tillering stage as well, but when we tried that during the pre-experiment, all plants died. The treatments were replicated six times in a randomized complete block design, and since our sampling was destructive, this resulted in 120 pots. After their 2-week exposure to drought, the plants were brought back to normal water content and not further exposed to drought. It was observed that during drought plants were weak but continued growth, that there was no delay in the occurrence of the next growth stage and that they showed resilience after rewetting. We sampled at tillering, stem elongation, booting, heading and flowering (Fig. 1). At stem elongation, booting and heading, we waited after the soil water content reached its target value (within a few days) before sampling all treatments and controls, to make sure we sampled during the drought event. Only the pots from the corresponding treatment were under drought at the time of sampling, while all other pots were well watered. All the plants from a pot were uprooted and firmly shaken, after which the soil remaining attached to the roots – the rhizosphere – was sampled, mixed thoroughly and snap frozen in liquid nitrogen. This resulted in 120 rhizosphere soil samples (3 treatments + 1 control x 5 growth stages x 6 blocks). Rhizosphere soils were transported to the lab on dry ice and then kept at -80℃ until extracted.

**Figure 1.**
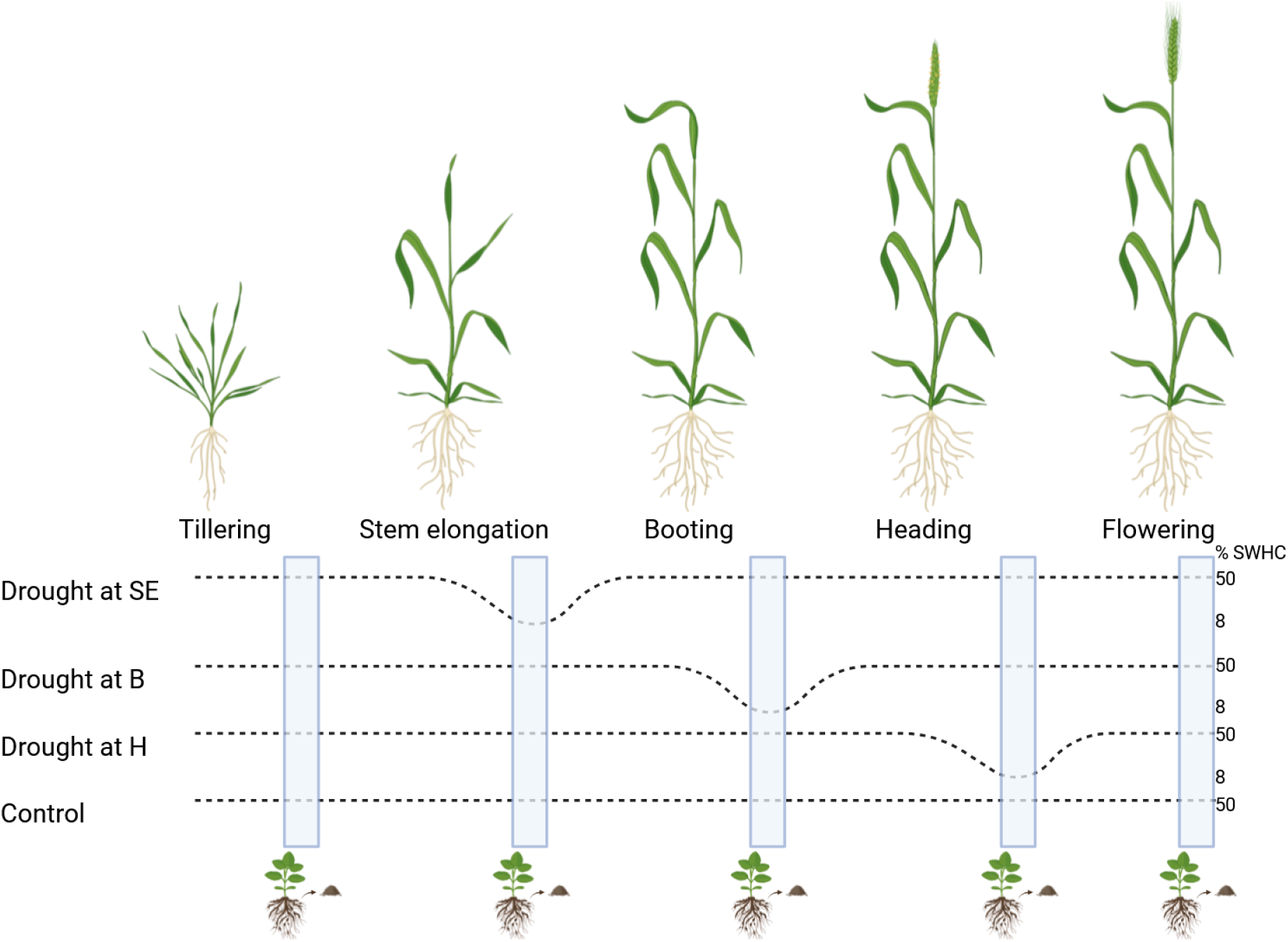
Experimental design. Water stress was imposed at Stem Elongation (SE), Booting (B) or Heading (H), and we sampled at Tillering, Stem Elongation, Booting, Heading, and Flowering. Created in https://BioRender.com

### RNA extraction and sequencing

Total RNA was extracted from 2 g of rhizosphere soil using the RNeasy PowerSoil Total RNA Kit (QIAGEN, Canada). Extracted RNA was treated with DNAse (ThermoFisher, Canada) to remove the DNA prior to sequencing. The absence of DNA was confirmed by the lack of PCR amplification using 16S rRNA gene specific primers. Total RNA was sent for two lanes of NovaSeq6000 S4 PE150 sequencing at the Centre d’Expertise et de Services Génome Québec (Montréal, Québec). Libraries were created using a microbial ribosome subtraction approach. The raw sequencing data produced in this study was deposited in the NCBI under Bioproject accession PRJNA978601.

### Bioinformatics analyses

The metatranscriptome sequencing of the 120 rhizosphere samples resulted in 12,000M reads for a total of 1,800 giga bases which were processed together through our metatranscriptomics bioinformatics pipeline [26], as detailed in Pande et al. [2]. For gene functional assignment, we used the JGI’s guidelines [27] including the assignment of KEGG orthologs (KO). For taxonomic assignment, each contig was blasted (BLASTn v2.6.0+) against NCBI’s nt database (version downloaded from NCBI’s server on January 9^th^ 2019) and the best hit’s taxonomic identifier was used to assign a taxonomic lineage to the contig. Taxonomic summaries were performed using MicrobiomeUtils v0.9 (github.com/microbiomeutils).

### Statistical analyses

All statistical analyses were performed in R version 4.2.3. [28]. Transcript differential abundance analyses between the drought treatments and the growth stages were carried out using the EBTest function of the EBSeq library with a false discovery rate (FDR) of 0.05. Anovas were performed using the aov function of the stats package. Permanovas were performed using the adonis2 function of the vegan package. For both Anova and Permanova, the blocks were considered in the formula. All graphs were generated using ggplot2. The R project folder containing the R code used for data manipulation, statistical analyses, and tables and figure generation is available on our lab GitHub repository (https://github.com/le-labo-yergeau/MT_Time_Wheat). The associated transcript abundance and annotation tables, and the metadata files used with the R code are available on Zenodo (https://doi.org/10.5281/zenodo.7995531).

## Results

### Growth stages have a strong influence on the wheat rhizosphere metatranscriptome

We performed permanova and principal coordinate analysis (PCoA) on the entire metatranscriptome (14,709,097 transcripts) to assess the effect of the four drought treatments and the five growth stages. 16.1% of the variation could be explained by growth stage (P=0.001), 3.2% by the drought treatments (P=0.008) and 11.1% by the interaction between the two variables (P=0.001). This can be visualized in the PCoA (Fig. 2a), where the clustering of the samples is mostly driven by the wheat growth stage, with some, less visible in the first two dimensions, effects of the drought treatments. We also looked at the treatments, but classified as when the drought was applied (DR), as rewetted (RW, treatments after the drought event) or as not disturbed (ND, controls and treatments before the drought event) (Fig. 2b). These results show that growth stage has a strong influence on the metatranscriptome.The drought treatments also affected the metatranscriptome, but its effect varied according to the wheat growth stage.

**Figure 2.**
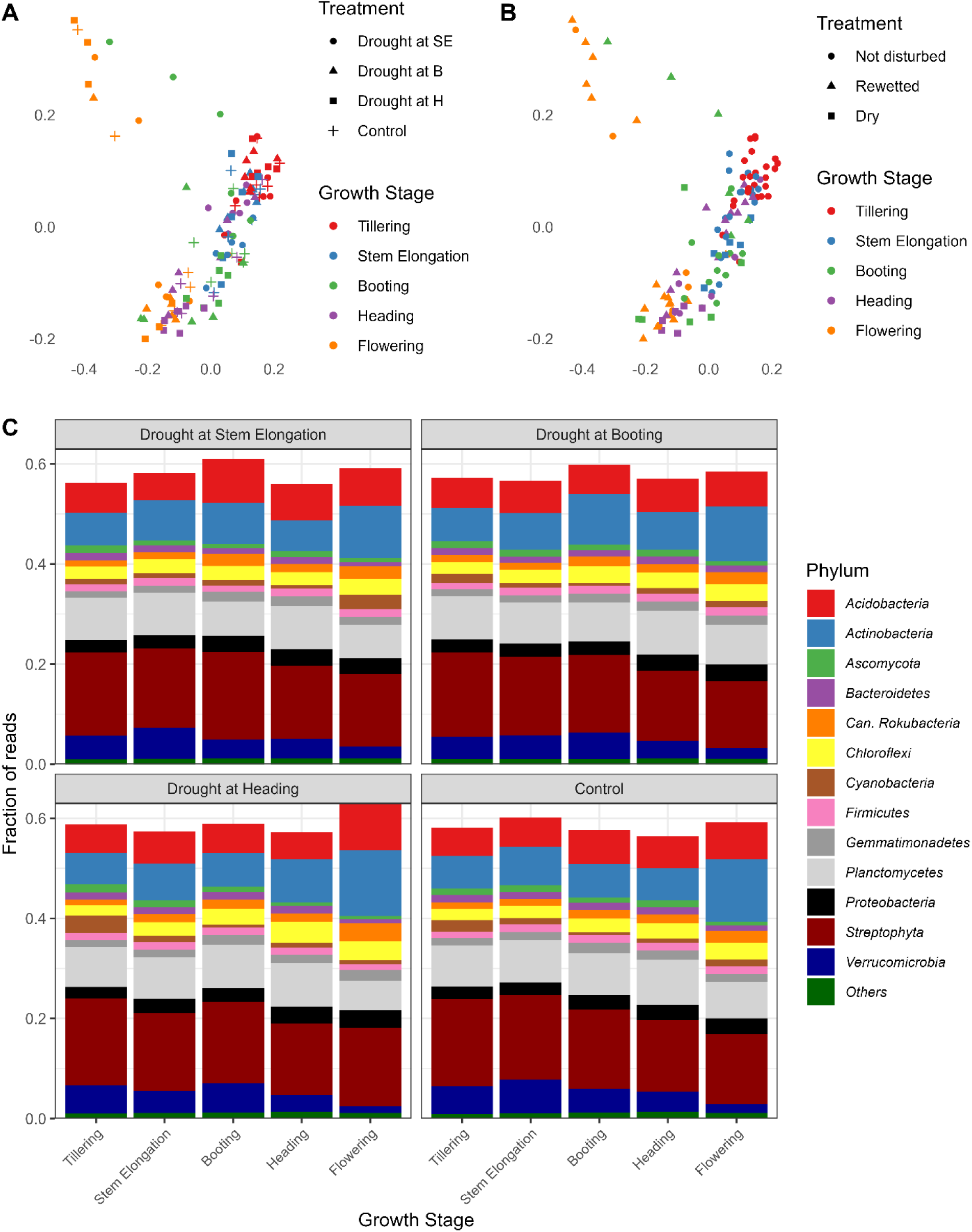
Growth stage has a strong influence on the wheat rhizosphere metatranscriptome. Principal coordinate analysis based on the relative abundance of all transcripts (A,B) and community composition at the phylum level (C) for the rhizosphere metatranscriptome of wheat plants at tillering, stem elongation, booting, heading and flowering stages that were stressed at stem elongation (SE), booting (B) or heading (H), or not stressed (control).

We then looked at the taxonomical affiliation of all transcripts across growth stages for all treatments (Fig. 2c) and tested the difference between the control and the different treatments during the drought or after rewetting using paired t-tests. Since there was quite some variability between the treatments, we also kept results with 0.05<P<0.10. As compared to the controls, the relative abundance of *Ascomycota* decreased 1.45 times during drought, when applied at stem elongation (t=2.49, P=0.062). When this treatment was rewetted, the *Bacteroidetes* were 1.34 times less abundant than in the control (t=2.31, P=0.069). When drought was applied at booting, none of the most abundant phyla changed their relative abundance, both during drying and rewetting. When soils were dried at heading, the *Actinobacteria* were 1.34 times more abundant than in the controls (t=4.95, P=0.004), whereas the *Chloroflexi* were 1.38 times more abundant (t=3.94, P=0.011). For the *Chloroflexi*, this difference was sustained during rewetting, where they were 1.14 times more abundant in the rewetted treatment as compared to the control (t=3.27, P=0.022).

We also looked at the COG category of the transcripts (Fig. 3), and tested differences using paired t-tests as above. As above, since there was quite some variability between the treatments, we also kept results with 0.05<P<0.10.When soils were dried at the stem elongation stage, transcripts related to “Carbohydrate transport and metabolism” were decreased by 1.10 times during drought as compared to the controls (t=2.49, P=0.055). When this treatment was rewetted, no difference was found with the controls. When drought was applied at booting, transcripts affiliated to “Energy production and conversion” and “Translation, ribosomal structure and biogenesis” decreased 1.12 and 1.16 times, respectively (t=2.23, P=0.077 and t=2.39, P=0.063, respectively), whereas transcripts classified in the “Transcription” category increased 1.15 times (t=2.06, P=0.095). No significant difference remained for this treatment after rewetting. When soils were dried at heading, transcripts classified in the COG categories “Posttranslational modification, protein turnover, chaperones” and “Transcription” increased 1.24 and 1.39 times as compared to the controls, respectively (t=2.26, P=0.074 and t=3.44, P=0.019, respectively). This difference was sustained at rewetting for the “Transcription” category that was 1.12 times more abundant than the controls (t=2.24, P= 0.075).

**Figure 3.**
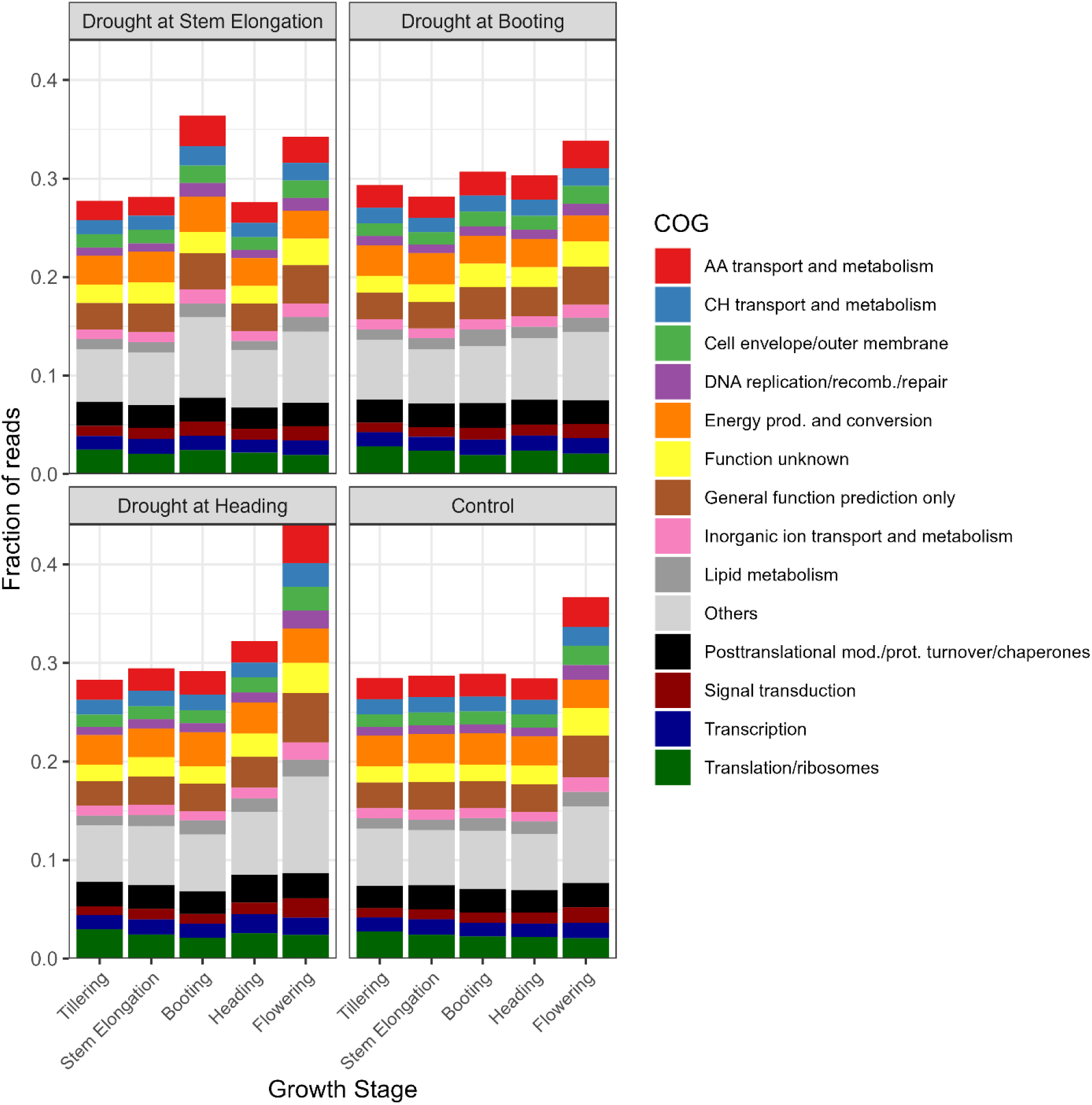
Growth stage also influences the functional composition of the metatranscriptome in the wheat rhizosphere. Community composition at the COG category level for the rhizosphere metatranscriptome of wheat plants at tillering, stem elongation, booting, heading and flowering stages that were stressed at stem elongation, booting or heading, or not stressed (control).

### The response to drought across growth stages is idiosyncratic

To analyze the effect of the drought treatments on the metatranscriptome, we did a transcript differential abundance (DA) analysis between the control and each of the three drought treatments across the different growth stages. In terms of number of differentially abundant transcripts, the patterns varied per treatment (Fig. 4a). When drought was applied at stem elongation, there was a six-fold increase in the number of DA transcripts from stem elongation to booting (from drought to rewetting), followed by a four-fold decrease at heading and another four-fold decrease at flowering (Fig. 4a). When drought was applied at booting, however, there was an 11-fold increase in the number of DA transcripts during drought (booting) followed by a sharp decrease at rewetting (heading) (Fig. 4a). Likewise, when drought was applied at heading, there was a four-fold increase in the number of DA transcripts during drought (heading) followed by a two-fold reduction at rewetting (flowering) (Fig. 4a). Taken together, these results show that the drought treatments resulted in large changes in the rhizosphere metatranscriptome. The amount of DA transcripts varied through the plant growth stages depending on the timing of the drought event.

**Figure 4.**
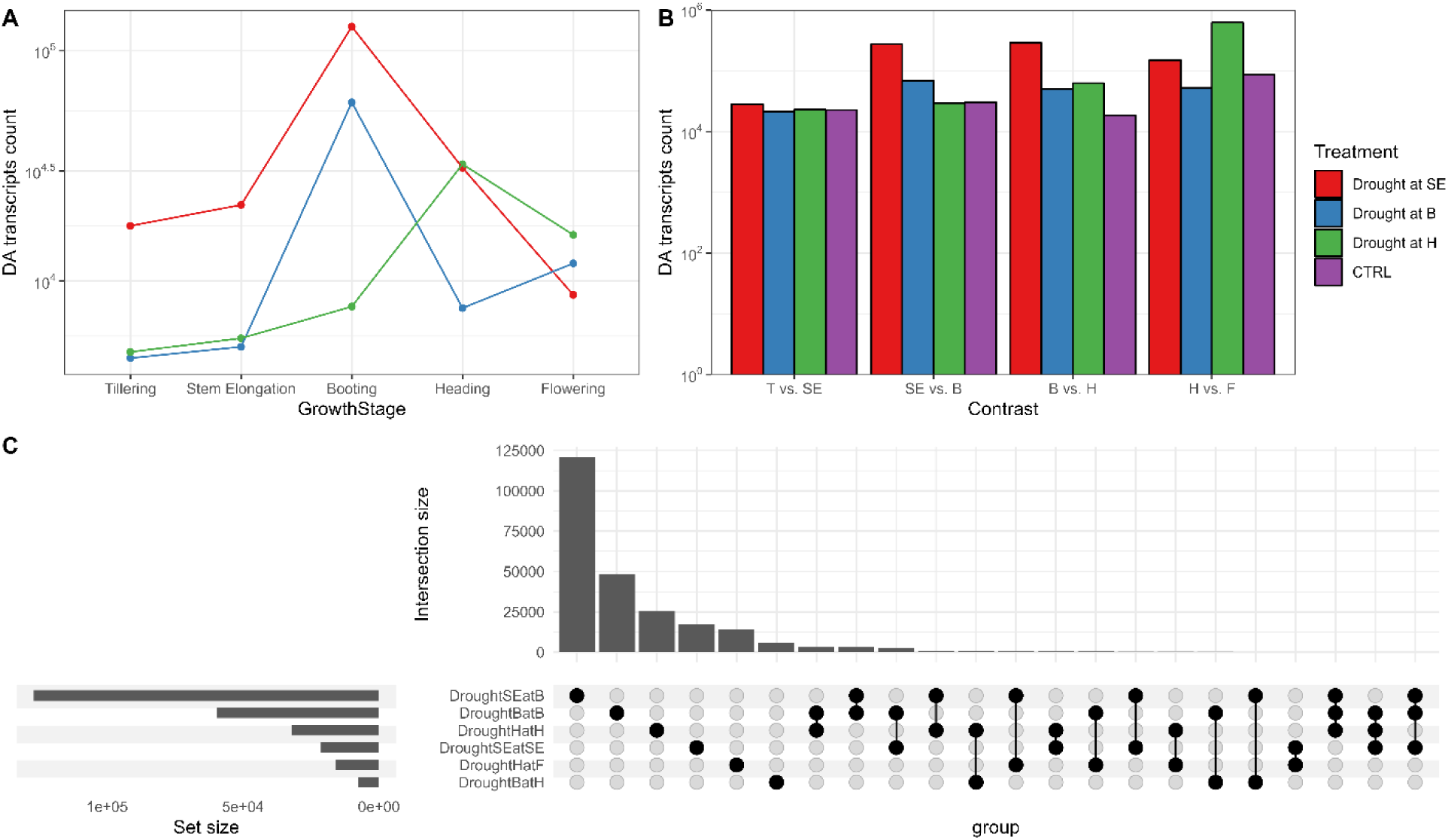
The response to drought across growth stages is idiosyncratic. Number of DA transcripts (treatment vs. control) in the rhizosphere metatranscriptome of wheat plants at tillering, stem elongation, booting, heading and flowering stages that were stressed at stem elongation, booting or heading (A). Number of DA transcripts between adjacent growth stages for the rhizosphere metatranscriptome of wheat plants that were stressed at stem elongation, booting or heading, or not stressed (B). Number of shared DA transcripts (treatment vs. control) in the rhizosphere metatranscriptome of wheat plants during drought at stem elongation, booting or heading, or after rewetting. Only intersection with size above 100 are shown (C). SE: stem elongation, B: booting, H: heading, F: flowering.

We did a co-occurrence analysis of the DA transcripts found at the dry and rewetting stages for the three treatments (Fig. 4c). Most of the transcripts were unique to a single treatment (Fig. 4c). For the drought at stem elongation treatment, 17,166/21,399 (80.2%) and 121,052/126,995 (95.3%) of the DA transcripts were unique at the stem elongation and booting stages, respectively. For the drought at booting treatments, 48,402/59,607 (81.2%) and 5,896/7,636 (77.2%) of the DA transcripts were unique at the booting and heading stages, respectively. Finally, for the drought at heading treatment, 25545/32141 (79.5%) and 14094/15877 (88.8%) of the DA transcripts were unique at the heading and flowering stages.

Across the six samples (total of 247,498 DA transcripts), the largest intersection was between the drought samples at booting and heading, but still only 3,512 DA transcripts were shared (Fig. 4c). 800 DA transcripts were shared by 3 or more treatment x growth stage combinations, 14 by 4 or more treatments x growth stage combinations, and none by 5 or more treatments x growth stage combinations (Fig. 4c). When comparing the samples taken during the drought events, 174 DA transcripts were shared by the three treatments, whereas the samples taken during the rewetting shared only 2 DA transcripts. Taken together, these results suggest that the response of the microbial communities to the drought treatment is very different when drought is applied at different plant growth stages, and that only a minority of DA transcripts are shared across the different treatments.

We then looked at the genus and COG function annotation of DA transcripts of Fig. 4a. A large portion (∼75%) of the DA transcripts could not be annotated at the genus and the COG function levels and were removed from the figure for clarity. As the results we are presenting are the taxonomic or the functional make-up of a list of DA transcripts generated by comparing all replicates, it is not possible to perform statistical comparisons as we end up with a single DA transcript list per treatment comparison. The taxonomic makeup of the DA transcripts varied across growth stages across all treatments, even before the drying and rewetting events (Fig. 5a). The total fraction of DA transcripts that belonged to the genera listed in Fig. 5a – that made up at least 0.25% of the DA transcripts across all samples – was lower at the stage that showed the largest increases in DA transcripts in Fig. 4a. This could be related to an increase in the unannotated DA transcripts or an increase in other genera that did not cross the inclusion threshold across all samples. Besides, there are no clear trends in the shifts among the genera that made up the DA transcripts. For the COG functions, however, the make-up of the DA transcripts was much less variable, with similar proportions of the most abundant functions across most stages for all treatments (Fig. 5b). There were some shifts visible, mostly at the stage where we had seen the largest number of DA transcripts (Fig. 4a), in many cases for COG functions related to amino acids (Fig. 5b). In any case, the shifts in the COG functions of the DA transcripts are subtler than the shifts observed at the genus level.

**Figure 5.**
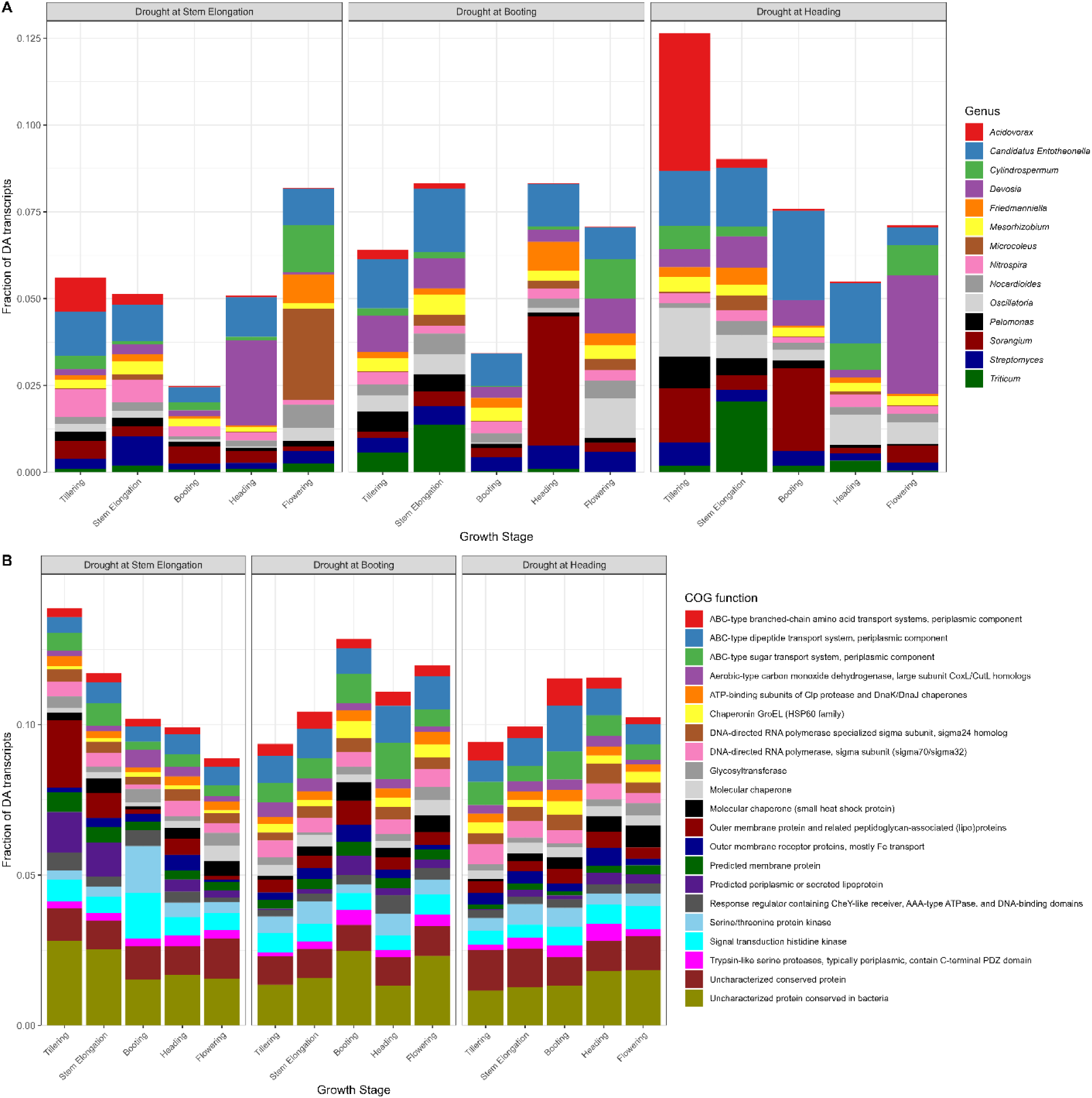
The taxonomical and functional makeup of the DA transcripts vary without clear patterns. Genus (A) and COG function (B) makeup of the DA transcripts (treatment vs. control) in the rhizosphere metatranscriptome of wheat plants at tillering, stem elongation, booting, heading and flowering stages that were stressed at stem elongation, booting or heading.

### Shifts in metatranscriptome across growth stages are compounded by drought

We also did differential transcript abundance analyses between each contiguous growth stage (tillering vs. stem elongation, stem elongation vs. booting, booting vs. heading, heading vs. flowering), for both treatments and control. We did this to disentangle the effects of growth stages on the metatranscriptome vs. the temporal effects of the drought treatments. Here again, since we incorporated all replicates in DA analyses, we end up with a single list of DA transcripts per treatment comparison, and it is therefore not possible to perform statistical comparisons. In the controls, the number of DA transcripts was 2.8 to 4.6 times higher for the heading vs. flowering contrast as compared to the other contrasts (Fig. 4b), suggesting large shifts in the rhizosphere metatranscriptome during flowering. For the drought at stem elongation treatment, there were many DA transcripts (150,083 to 287,091) for the stem elongation vs. booting, booting vs. heading and heading vs. flowering contrasts, which was up to 15.5 times more than the controls (for the booting vs. heading contrast) (Fig. 4b). For the drought at booting treatment, the number of DA transcripts was higher than in the controls only for the stem elongation vs. booting and booting vs. heading contrasts (Fig. 4b), up to a factor of 2.7. For the drought at heading treatment, the number of DA transcripts was higher than the controls for the booting vs. heading and the heading vs. flowering contrasts (Fig. 4b), with a 7.3 time increase for the latter case. These results indicate that the microbial metatranscriptome does change with plant growth stages, but that these changes are compounded by water stress.

Most of the DA transcripts were not shared between the treatments, for all contrasts including the ones before drying (Fig. 6). This suggests that at the transcript level, there was a lack of a coherent temporal response of the rhizosphere microbiome to drought and to growth stage. The genus-level affiliation of the DA transcripts varied across the contrasts (Fig. 7a).

**Figure 6.**
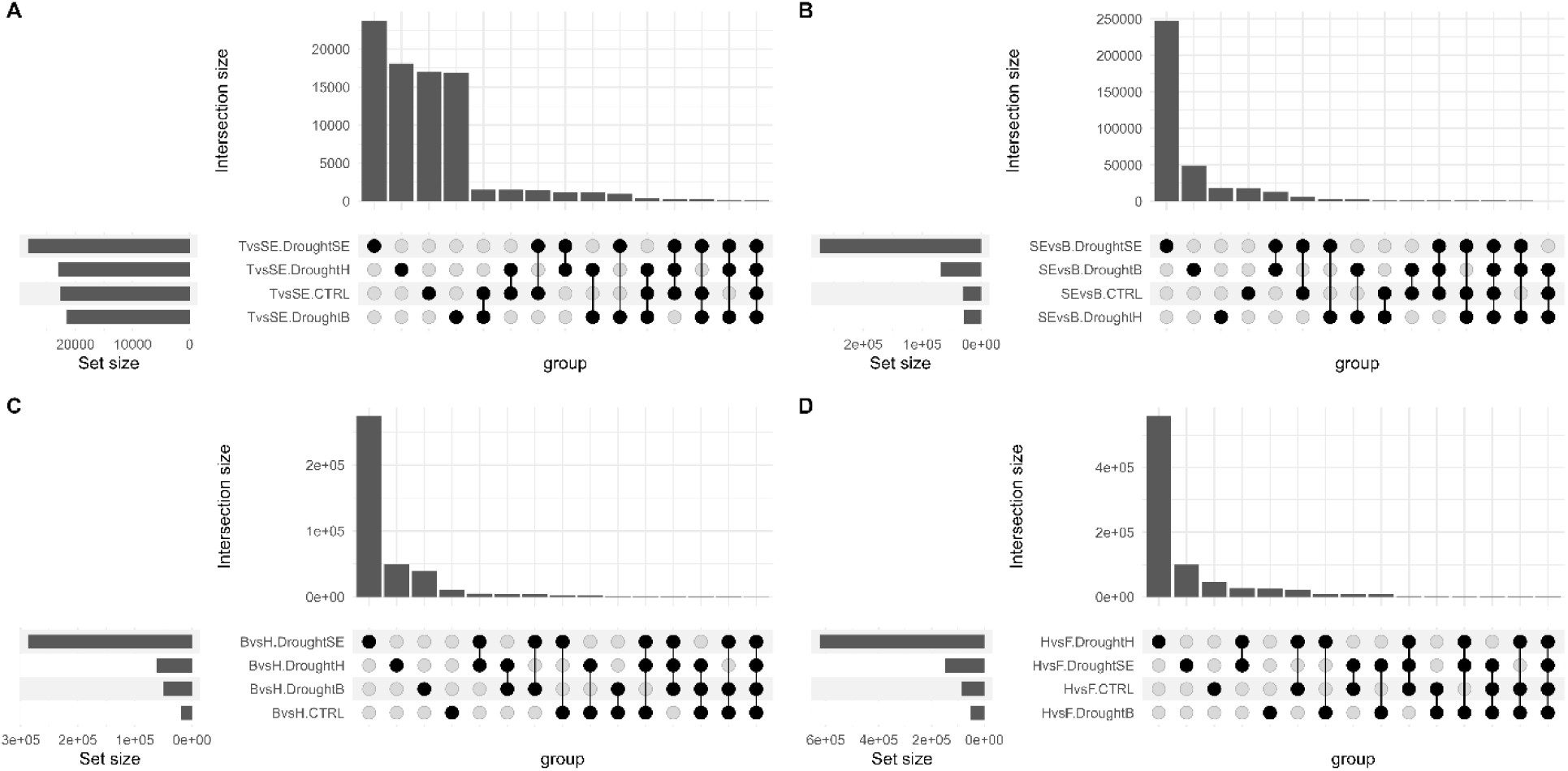
Drought treatments affect the normal microbial succession across growth stages. Number of shared DA transcripts between tillering and stem elongation (A), stem elongation and booting (B), booting and heading (C), and heading and flowering (D) for the rhizosphere metatranscriptome of wheat plants that were stressed at stem elongation, booting or heading, or not stressed. T: tillering, SE: stem elongation, B: booting, H: heading, F: flowering.

**Figure 7.**
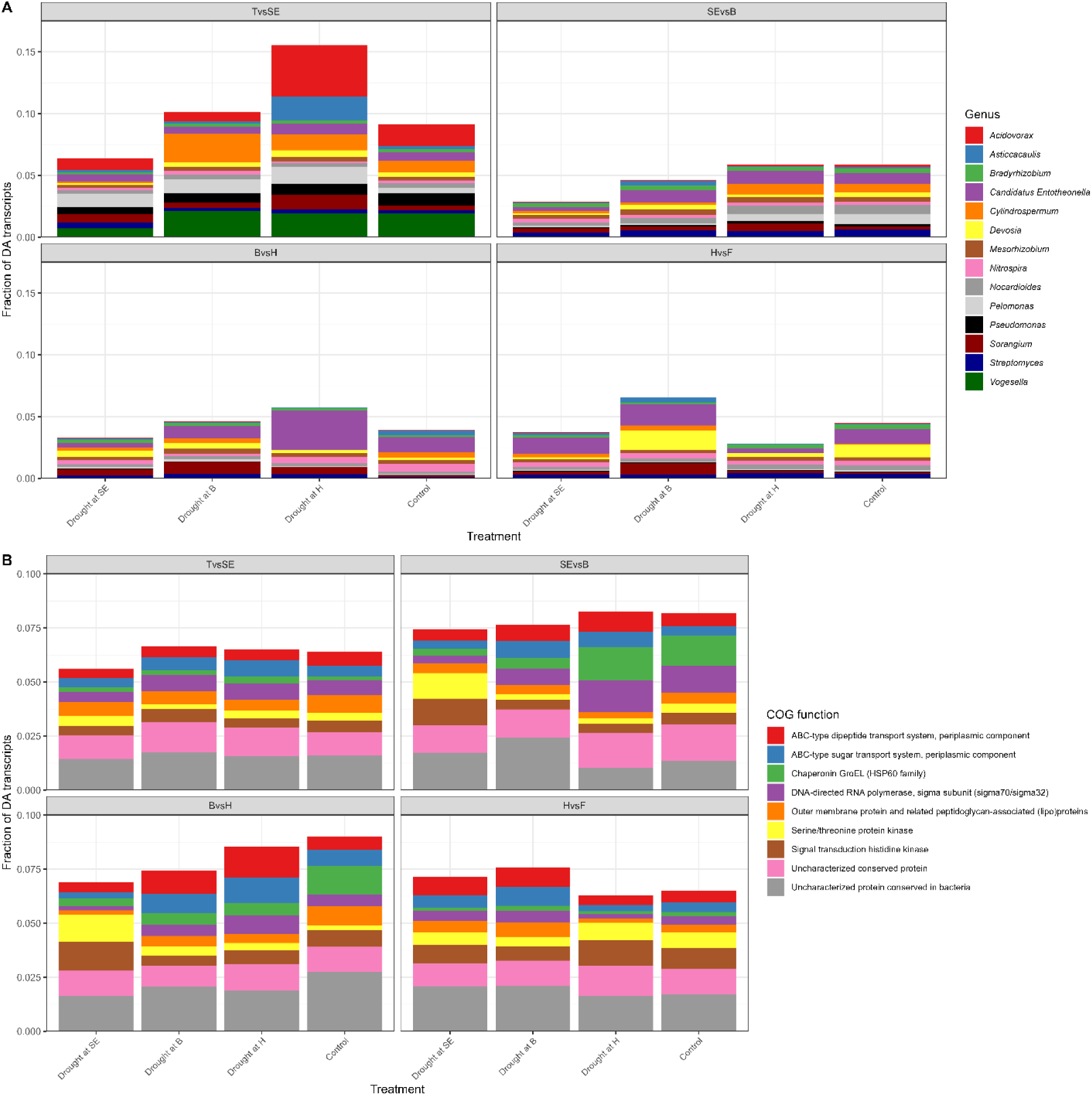
Drought treatments affect the normal microbial taxonomical and functional succession across growth stages. Genus (A) and COG function (B) makeup of the DA transcripts between adjacent growth stages for the rhizosphere metatranscriptome of wheat plants that were stressed at stem elongation, booting or heading, or not stressed. T: tillering, SE: stem elongation, B: booting, H: heading, F: flowering.

Although there were some common taxa represented in the DA transcripts, the water-stressed microbial succession differed from the well-watered (control) microbial succession (Fig. 7a). For instance, when comparing the stem elongation vs. the booting stages, the DA transcripts for the drought at stem elongation and booting treatments contained less *Cylindrospermum* and *Pelomonas* genera as compared to the controls (Fig. 7a). When comparing the booting and heading stages, the DA transcripts in all the treatments were enriched in *Sorangium* and depleted in *Cylindrospermum* and *Nitrospira* (Fig. 7a). Additionally, the drought at heading treatments transcripts were enriched in *Entotheonella*, whereas it was the inverse for the two other treatments. (Fig. 7a). For the heading vs. flowering contrast, the DA transcripts of the drought at booting treatments were more often affiliated to *Asticcacaulis*, *Devosia*, *Cylindrospermum* and *Sorangium* than the control DA transcripts (Fig. 7a). In contrast, the two other treatments showed inverse trends.

At the functional level – COG functions – the affiliation of the dominant DA transcripts was relatively stable for the control treatment (Fig. 7b). For the drought at stem elongation treatment, there were more DA transcripts identified to “Serine/threonine protein kinase” and “Signal transduction histidine kinase” than in the control for both the stem elongation vs. booting and booting vs. heading contrasts (Fig. 7b). For the stem elongation vs. booting contrast, there were less DA transcripts related to “Chaperonin GroEL” and “DNA-directed RNA polymerase” in the drought at stem elongation and at booting treatments, relative to the control (Fig. 7b). For the booting vs. heading contrast, the DA transcripts identified in the three treatments were depleted in “Chaperonin GroEL” and “Outer membrane membrane protein…” as compared to the control (Fig. 7b). Both the drought at booting and drought at heading treatments had more DA transcripts related to “ABC-type dipeptide transport system” and “ABC-type sugar transport system”. Finally, for the heading vs. flowering contrast, the affiliations of the DA transcripts of the treatments were like the control (Fig. 7b). There was, however, more DA transcripts in the “ABC-type dipeptide transport system” and “ABC-type sugar transport system” and “Outer membrane protein…” for the drought at stem elongation and drought at booting treatments, relative to the control (Fig. 7b). Taken together, these results suggest that at the genus and COG function levels, there are some groups that are consistently showing a difference in their succession when subjected to drying and rewetting.

## Discussion

We had previously shown that, within the wheat holobiont, the strongest responders were the microbial communities [2]. Here, we looked at how these microbial communities respond to a 2-week dry spell imposed at different growth stages. For all growth stages, drying resulted in many differentially abundant (DA) transcripts. However, only a small percentage of these DA transcripts were shared across growth stages, suggesting idiosyncratic responses. This response altered the microbial succession across wheat growth stages.

When we imposed drought at an early stage (tillering), we killed all our wheat plants. Similarly, many plants resist better to disease as they age [29]. Plant aging also modulates the response to drought of the associated microbial communities. For example, the root microbial community composition of older sorghum plants is less impacted by drought than the one of younger plants [30]. Our youngest plants that survived drought (stem elongation stage) were the ones that showed the largest metatranscriptomic shifts. These shifts were larger, more intense during rewetting and lasted longer, as compared to the ones observed in older plants. For these older plants – at the booting and heading stages – the metatranscriptome showed smaller changes that rapidly returned to unstressed control levels. It therefore appears that, in agreement with our hypothesis, the rhizosphere metatranscriptome of younger wheat plants reacts more to drought.

On top of having different magnitude, the shifts observed at the transcript level across the drought treatments were idiosyncratic. Idiosyncrasy is often seen in the response of microbial communities to various factors. For instance, depending on the plant compartment and the water stress history of the soil, drought altered differently the wheat microbial communities [13–15]. The wheat microbial community is also influenced by wheat genotype, but constrained by the plant growth stages, compartment and time [16]. This idiosyncrasy also depends on the resolution of the analyses: at coarse taxonomic or functional levels, some patterns are conserved (e.g. increased relative abundance of *Actinobacteria* following drought) [13, 15, 31–33], but at finer levels, there is no coherent response. For instance, the microbial responses to drought at two different plant growth stages differed at the OTU level but resulted in the same enrichment of monoderms [30]. Here, most of the DA transcripts were unique for a single treatment, but when looking at functional categories (i.e. COG functions), the differences were less important. This supports the idea that shifts at fine taxonomic or at the individual transcript levels might be seemingly random but are coherent at the functional category level or at coarser taxonomical levels [31, 34]. Alternatively, part of the idiosyncratic responses observed could be due to stochastic variation, as even before the drought treatments were applied, many of the DA transcripts were unique and showed distinct taxonomic affiliations across treatments.

Some of the idiosyncrasies observed could be explained by the microbial succession through plant developmental stages. As plant develops, its microbial community fluctuates, and drought does not affect similarly microbial communities of different diversity and composition. For instance, Gram-negative bacteria produce osmolytes in response to drought, whereas they are constitutively produced in Gram-positive [35]. At the same time, Gram-positive bacteria generally have a thicker cell wall that Gram-negative bacteria [36]. Accordingly, Gram-positive bacteria (mostly *Actinobacteria* and *Firmicutes*) made up 90% of the wheat microbial isolates that could grow under high osmotic pressure [37]. As mentioned above, *Actinobacteria* generally increase in relative abundance following drought [13, 15, 31–33], suggesting a heightened capacity to resist to that stress. The relative activity of *Actinobacteria* also increased at the flowering stage in our study. As compared to bacteria, the wheat fungal community composition often doesn’t respond as much to changes in soil water content [17, 38], but they strongly responded to decreasing soil water content at the metatranscriptomic level [2] and many fungal isolates from wheat could grow at high osmotic pressure [37]. Hence, the magnitude of the metatranscriptomic response of the microbial community to drought will likely be defined by the relative dominance of fungi, Gram-positive bacteria, and Gram-negative bacteria, which all have different life-strategies when facing water stress [31, 38, 39]. As all these groups vary through plant development, this could partly explain the contrasting responses of the microbial communities to drought imposed at different plant growth stages.

In addition to its strong idiosyncratic effect on microbial transcripts, drought resulted in a different microbial transcriptomic succession as compared to the well-watered controls. Previous studies had shown that the plant root exudates varies across plant growth stages [20], shifting the microbial community and the metatranscriptome of the rhizosphere [16, 21, 40, 41]. The effect of drought on microbial communities was, however, shown to trump the effect of plant development [30]. Indeed, across developmental stages, we found a much larger number of DA transcripts in the treated rhizospheres as compared to the controls. In the controls, the largest number of DA transcripts was found when comparing heading to flowering. The flowering vs. heading comparison was also the one that yielded the largest number of DA transcripts in the treated samples, probably because it compounded the large shifts due to succession with the ones due to drought. Under drought, the microbial transcriptomic succession in the rhizosphere was heightened and had little to do with the well-watered (control) succession, even though the development of the plants was not visibly disrupted by the drought.

We worked with a single wheat genotype that was not bred for resistance to water stress. The response of wheat to water stress does vary across genotype, with some being better at tolerating water stress at particular growth stages [42]. The wheat microbiome also varies across genotypes [13, 15], but this is constrained by plant compartment, growth stages and with environmental conditions [16]. Historical exposure of the soil microbiome to water stress also modulates its interaction with wheat genotypes [13–15]. It would be important to confirm the results presented here for genotypes with a gradient of water stress resistance and across multiple soils, ideally in the field.

We showed here that drought strongly affects the wheat rhizosphere microbial transcriptome, but in a growth stage-dependent fashion. Our results imply that - as for the plant - early drought events impact more the microbial communities. These altered microbial communities might be less beneficial, which could partly explain the vulnerability of young plants to stresses. Microbial communities also responded when drought was applied at other growth stages, but the identity of the transcripts was very different. In all cases, drought altered the microbial transcriptome of the rhizosphere, which could have repercussions on wheat growth and grain quality. Understanding the interaction between crop growth stage and drought events on the rhizosphere microbiome is essential to optimize crop resistance and resilience under the current climate emergency.

## Data availability

The raw data produced in this study was deposited in the NCBI under Bioproject accession PRJNA978601. The R project folder containing the R code used for data manipulation, statistical analyses, and tables and figure generation is available on our lab GitHub repository (https://github.com/le-labo-yergeau/MT_Time_Wheat/). The associated transcript abundance and annotation tables, and the metadata files used with the R code are available in the Zenodo archive (https://doi.org/10.5281/zenodo.7995531).

## Author contributions

PMP: Designed, set-up and maintained the greenhouse experiment, performed the soil sampling and molecular analyses. JT: Performed the bioinformatic analyses. MSA: Designed the experiment, supervised students and secured funding. EY: Designed the experiment, performed the statistical analyses, secured funding, supervised students and wrote the manuscript with inputs from all authors.

